# Respiration shapes response speed and accuracy with a systematic time lag

**DOI:** 10.1101/2024.09.09.611983

**Authors:** Cosima Harting, Lena Hehemann, Lisa Stetza, Christoph Kayser

**Affiliations:** Department for Cognitive Neuroscience, Faculty of Biology, Bielefeld University, Bielefeld, Germany

**Keywords:** perception, cognition, respiration, reaction times, performance

## Abstract

Sensory-cognitive functions are intertwined with physiological processes such as the heart beat or respiration. For example, we tend to align our respiratory cycle to expected events or actions. This happens during sports but also in computer-based tasks and systematically structures respiratory phase around relevant events. However, studies also show that trial-by-trial variations in respiratory phase shape brain activity and the speed or accuracy of individual responses. We show that both phenomena, the alignment of respiration to expected events and the explanatory power of the respiratory phase on behaviour co-exist. In fact, both the average respiratory phase of an individual relative to the experimental trials and trial-to-trial variations in respiratory phase hold significant predictive power on behavioural performance, in particular for reaction times. This co-modulation of respiration and behaviour emerges regardless of whether an individual generally breathes faster or slower and is strongest for the respiratory phase about two seconds prior to participant’s responses. The persistence of these effects across 12 datasets with 277 participants performing sensory-cognitive tasks confirm the robustness of these results, and suggest a profound and time-lagged influence of structured respiration on sensory-motor responses.

## Introduction

Recent studies highlight the relation between physiological processes such as respiration, heartbeat, gut activity and brain function [1-4]. Taking respiration as an example, we know that the brain structures controlling respiration and those sensing the resulting changes in airflow or chest pressure are intricately connected with the limbic and autonomous nervous system [5-7]. As a result, direct neural feedback about the current respiratory state is widely available in the brain [8-10]. In addition, respiration-driven changes in blood oxygenation and arterial perfusion affect the brain’s metabolism and its capacity to supply neural processes with energy. Hence, respiration can both directly and indirectly shape how sensory information is encoded and translated into cognitive decisions or motor actions [11-13].

Several studies show that participants’ trial-to-trial performance in sensory-cognitive tasks covaries with the respiratory phase [12, 14-19] [11]. Still, it remains unclear whether this co-modulation of behaviour with respiration relates to how good individuals perform a task (perceptual or cognitive accuracy), how fast they react (reaction times), or both. Reaction times may vary with respiration simply due to changes in muscle tension or brain-muscle coordination with the respiratory cycle [20-23]. In contrast, a covariation of response accuracy or perceptual sensitivity along the respiratory cycle may result from changes in neural excitability or inter-areal coupling [13, 24], and hence point to a more profound modulation of sensory-cognitive processes. Furthermore, most studies focused on the immediate respiratory phase in each trial. However, given the various pathways by which respiration-related signals or changes in blood oxygenation can effect brain function, it remains unclear whether any co-modulation of respiration and behaviour is immediate or temporally delayed [5].

At the same time, humans tend to align their respiration to expected events or actions. For example, holding breath is often used during shooting and fast motor actions in sports are accompanied by specific breath patterns [25, 26]. Similarly, the respiratory pattern of participants during neuro-cognitive laboratory tasks is structured and exhibits consistency across trials [27, 28]. However, perfect consistency of the respiratory phase across repeats of the same action would leave little trial-to-trial variability in respiration to explain variability in the action outcome. We here investigate how both a systematic alignment of respiration to task-events and trial-to-trial variations in respiratory phase that is predictive of behaviour can co-exist.

We leverage a large dataset (data from 277 individuals) to systematically probe the alignment of respiration to experimental tasks and the relation between task performance (response accuracy, reaction times) and respiratory phase. This dataset comprises 12 experiments implementing sensory and cognitive tasks (of these, 6 published previously but re-analysed here). Participants performed these tasks without a specific constraint on how to breathe. This instruction was used to mimic typical ‘every-day’ experiments that do not impose specific manipulations on participant’s respiration. The results show that participants tend to align their respiration to individual trials, resulting in a consistent pattern of respiration both across trials for individual participants and on average across trials. At the same time behavioural performance is co-modulated with the residual variability of the respiratory phase. This co-modulation is strongest when measured against the respiratory phase about 2 seconds prior to the individual responses.

## Methods

### Participants

All studies were approved by the ethics committee of Bielefeld University. Adult volunteers participated after providing informed consent and were compensated for their time. All had self-reported normal vision and hearing and none indicated a history of neurological disorders. The data were collected anonymously and it is possible that some individuals participated in more than one of the experiments described below. Demographical data were not collected, but we expect these to be very similar to previous studies, in particular as the participant pool consisted of typical young university students [29, 30]. The specific interest to investigate the relation between respiration and task performance was not mentioned explicitly prior to the study. Participants were instructed to “breathe through their nose as usual”, as if performing the experiments without wearing the mask. During the experiment we could not continuously monitor whether participants adhered to this instruction, leaving the possibility that during parts of the experiment participants were breathing orally.

### General procedures

The experiments were performed in a darkened and sound-proof booth (E: Box; Desone, Germany). In one line of experiments, visual stimuli were presented on a computer monitor (27” monitor; ASUS PG279Q, about 1m from participants head) while acoustic stimuli were presented from two speakers placed besides the monitor. In another line of experiments participants sat in front of an acoustically transparent screen (Screen International Modigliani, 2×1 m, about 1m from the participants head) onto which visual stimuli were projected (LG HF65, LG electronics) while acoustic stimuli were presented from headphones (Sennheiser DH200Pro). Acoustic stimuli were calibrated to have an r.m.s. level of about 65 dB SPL. Stimulus presentation was controlled using the Psychophysics Toolbox (Version 3.0.14) using MATLAB (Version R2017a; The MathWorks, Inc., Natick, MA) and was synchronized to a BioSemi EEG recording system using TTL pulses. Participants responded using computer keyboards.

### Recording of respiratory signals

Respiration was recorded using a temperature-sensitive resistor that was inserted into disposable clinical oxygen masks (Littelfuse Thermistor No. GT102B1K, Mouser electronics). This effectively captures the continuous temperature changes resulting from the respiration-related airflow [27, 31]. The voltage drop across the thermistor was recorded via the analogue input of an ActiveTwo EEG system (BioSemi BV; Netherlands) at a sampling rate of 500 or 1000 Hz. We verified that the voltage drop of the temperature sensor follows the respiratory air-flow without time lag. For this we combined the temperature probe with two short-latency airflow sensors (F1031V, Mass Airflow Sensor, Winsen) and confirmed that the temperature change tightly aligns with the directional change in airflow.

### Behavioural paradigms

We analysed both data from experiments collected de novo (datasets 1-7) and from experiments performed and published in a previous study (datasets 8-12)[27]. All experiments involved one or more blocks of experimental trials, a practice block and possibly a block to determine each participant’s perceptual threshold for the respective task. Respiratory data were only collected during the main experiments. For each task, participants were instructed to respond as fast and accurately as possible after stimulus presentation.

Trials started with a fixation period (400-1000ms, uniform; unless stated otherwise) and inter-trial intervals were 1200-1500ms (uniform).

#### Datasets collected in this study

Pitch discrimination 1: Participants had to compare the pitch of two brief successive tones, as used in previous studies [32]. During each trial two pure tones (50ms duration, 6ms cosine ramp, 50ms pause in between, 65 dB SPL) were presented and participants had to indicate which of the two (first, or second) had higher pitch. Pitch differences were titrated around each participant’s psychophysical threshold in five levels, with a total of 600 trials. Data from N=16 participants.

Time perception: Participants had to categorize the duration of acoustically presented intervals as either ‘short’ or ‘long’. Intervals lasted 200, 250, 300, 350 or 400ms, with the 300ms ms being ambiguous and trials not analysed. Stimuli were 1024Hz pure tones and participants performed 540 trials. Data from N=18 participants.

Emotion discrimination 1: Participants had to discriminate the emotional expression (sad or happy) in briefly presented faces (subtending about 15 × 12 degrees, presented for ∼17 ms). Stimuli were obtained from the Dynamic FACES database [33]. For 24 individual faces we selected images that either reflect each emotion clearly, each emotion to a mild degree or were neutral. Hence, emotions were expressed in 5 levels, with the intermediate level being emotionally neutral and not analysed here. Each participant completed 480 trials. Data from N=25 participants.

Visual shape discrimination 1: Participants had to discriminate the orientation of briefly presented Necker cubes (subtending about 11 × 9 degrees, presented for ∼50ms). For each cube four edges had an increased luminance so to render the orientation of the shape less ambiguous, with a total of 5 levels (with the intermediate level being ambiguous and not analysed here). Each participant completed 480 trials. Inter-trial intervals were 800-1200ms (uniform), fixation periods were 1100-1500ms. Data from N=25 participants.

Pitch discrimination 2: This was the same as in pitch discrimination 1, except that there were only two levels of pitch difference titrated around each participant’s threshold. Prior to each block of this task participants performed one minute of structured respiration training (either box-breathing or hyperventilation). For the present study we collapse the data across both types of training. 480 trials per participant. Data from N=52 participants.

Emotion discrimination 2: Participants had to discriminate the emotion (anger or disgust) in briefly presented faces (subtending about 10 × 10 degrees, presented for 128 ms). The images were taken from the FACES database [34]. A total of 100 images was used and each participant completed 400 trials. Prior to each block of this task participants performed one minute of structured respiration training (either box-breathing or hyperventilation). For the present study we collapsed the data across both types of training and obtained 523 trials per participant. Data from N=24 participants.

Arithmetic: Participants performed arithmetical tasks requiring the summation of either a one- and a two-digit number (easy trials) for two two-digit numbers (difficult trials). Participants were presented the respective numbers on the screen and were given a maximal of 10 s to enter the respective sum. Participants performed a total of 270 trials, of which 135 were considered easy and 135 as difficult. Prior to each block participants performed one minute of structured respiration training or breathed normally. For the present study we collapsed the data across all types of respiration. Data from N=18 participants.

#### Previously published datasets

These have been described in detail in [27]. From this previous publication we re-analysed all five sensory tasks (ignoring the memory paradigm also reported there) using revised procedures.

Visual random-dot motion: Participants judged the direction of motion (left- or right-wards) of visual random dot displays lasting 340ms. The motion coherence varied across five levels around participant’s individual thresholds. Data from N=18 participants.

Pitch discrimination 3: This experiment is identical to pitch discrimination 1, with 400 trials per participant. Data from N=20 participants.

Pitch discrimination 4: This was similar to the above, except that participants had to manually initialize the start of the trial by pressing a key, which was then followed by a shorter fixation period (300 to 600ms uniform). Each participant completed 200 trials. Data from N=20 participants.

Sound detection: Participants had to detect acoustic sound (100 ms 1024 Hz pure tone) in a white noise background. Target came at one of four levels spaced around participant-specific thresholds or were absent. Each participant completed 480 trials. Data from N=20 participants.

Emotion discrimination 3: Participants had to discriminate the emotion (anger or disgust) in briefly presented faces (subtending about 8 degrees, presented for ∼100 ms). The images were taken from the FACES database [34]. A total of 400 images was used and each participant completed 400 trials. Data from N=21 participants.

### Analysis of respiratory data

Compared to our previous work [27] we improved the processing pipeline for respiratory data. The respiratory signals were filtered using 3-rd order Butterworth filters (high pass at 0.03 Hz, low pass at 6 Hz) and subsequently resampled at 100Hz using the FieldTrip toolbox [35]. To detect individual respiratory cycles, we applied the Hilbert transform and determined local peaks based on the respective phase [36]. Individual respiratory cycles were determined based on the data in windows of 7 seconds around each peak, whereby peaks were included only if the z-scored trace exceeded z=0.5 [12]. The inspiration period was defined as the continuous period with positive slope prior to the local peak (whereby interruptions of the positive slope shorter than 500ms were interpolated). The expiration period was defined as the continuous period with negative slope subsequent to the local peak (again interruptions shorter than 500ms were interpolated). This definition effectively splits the respiratory cycle effectively into the two main periods of inhalation and exhalation; though for some cycles short exhale pauses were classified as third state and not analysed [36]. In particular, compared to previous work [27] this procedure assigned a defined inspiration/expiration phase to more time points than in the previous work. To characterize atypical respiratory cycles, we compared the overall time courses of individual respiratory cycles using their mean-squared distances. We calculated the participant-wise distributions and excluded cycles with a distance larger than 3 standard deviations from the centroid as atypical. These cycles were excluded as they do not reflect the prototypical respiration under investigation here. Further individual trials were excluded from analysis as noted below. From the full datasets we retained only participants for which these procedures excluded less than 30% of the available trials for the final statistical analysis.

To link respiratory signals to behaviour we defined the phase of each respiratory cycle as a linearly increasing variable from the beginning to the end of inspiration (defined as angle from 0 to pi) and subsequently as linearly increasing from the beginning to the end of expiration (defined as pi to 2*pi). This phase variable scales linearly in time within each inspiration or expiration period and has two time points (0/2pi, and pi) whose interpretation is consistent regardless of the shape of individual respiratory cycles.

### Statistical analysis of the alignment between respiration and paradigm

To probe whether and how participants aligned their respiratory behaviour to the experimental trials we computed the phase consistency (also known as phase-locking) across trials. This was done by converting the phase into a complex-valued number, averaging this across trials and taking the vector length. To test for the statistical significance of this, we derived a surrogate distribution of phase-locking values under the null hypothesis of no alignment between respiratory trace and paradigm for each participant. This was obtained by randomly time-shifting the respiratory trace and recalculating the phase consistency 4000 times. We then compared the actual group-median distribution of group-medians in the surrogate data, taking the maximal value long the time axis to correct for multiple comparisons. We also tested whether this alignment of respiration to the experimental trials differed between trials with fast/slow responses, correct/wrong responses, and trials in the first/second half of each experiment. For reaction times we relied on a median split of trials, while for time-on-the-experiment we split trials based on the actual trial numbers. For response accuracy the number of correct and wrong trials was highly imbalanced. As this may bias the phase-locking estimate, we used equal trial numbers for each group, and excluded participants for which less than 40 trials were available per group.

### Analysis of the co-modulation of behaviour with respiration

For this we relied on linear mixed effect models (eq. 1). Separate models were fit for reaction times and accuracy, for each of the datasets, and separately for the phase of respiration at different time points relative to the experimental paradigm. As predictors we included a variable reflecting a parametric manipulation of task difficulty (e.g. the pitch difference), or for those tasks without such a parametric variable, one reflecting the two conditions to be discriminated. We also included a participant-wise random effect of trial number to capture potential effects of fatigue or training-on-the-task and a random offset for each participant. Respiration was modelled using the sine and cosine transformed trial-wise phase of respiration, at a particular time point of interest. These time points were either the time of stimulus onset, the time at which participants responded in each trial, or a time point at a particular lag preceding either stimulus onset or responses (defined in steps of 300ms). Note that this model implies a consistent relation between behaviour and respiratory phase across participants, if such an effect exists.

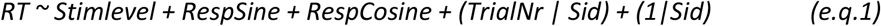

Reaction times were square-root transformed before entering the model. Models for reaction times used a linear activation function and assumed Gaussian variables, models for accuracy assumed binomial variables and a logistic activation function.

### Statistical analysis

We derived two measures of statistical significance for an effect of respiration in these models. The first focused on the predictive power of respiration to explain trial-to-trial variability in behaviour. For this we compared the Akaike information criteria (AIC) between two models, of which only one included respiration as factor. Based on this we derived the conditional probability of each model given the data, i.e. the associated Akaike weights [37]. We considered an AIC difference > 9.2 as evidence for an effect of respiration, which corresponds to the probability of the model including respiration to have more explanatory power than the alternative model of above 99%.

As a second measure of significance we focused on the slope of the respiratory predictors, which reflects the mean relation between respiration and behaviour. For this we compared the vector strength of the combined sine and cosine respiration predictors in the actual data to those in surrogate data [9, 27]. To obtain surrogate data we shuffled the trial-wise relation of the dependent and independent variables and computed the models again, repeating this procedure 4000 times. We then derived the p-value of the actual vector strength against the surrogate data for each experiment. The resulting 12 p-values were then combined across datasets using Fisher’s procedure [38]. We implemented this statistical test using the respiratory phase at three individual time points (−2.5s, 0s, +2.5s) relative to the stimulus or response (Table 1).

**Table 1.** Significance of the combined respiratory predictors. We derived the significance of the vector length of the sine- and cosine-respiration predictors for each dataset using a randomization test and combined the 12-resulting p-values across datasets using Fishers’ procedure. For this combination we report the associated t- and p-values. This analysis was performed at three time points of respiration relative to either the stimulus or response (−2.5, 0 and +2.5s).

|  | Stimulus aligned |  |  | Response aligned |  |  |
| --- | --- | --- | --- | --- | --- | --- |
|  | -2.5s | 0s | +2.5s | -2.5s | 0s | +2.5s |
| **Accuracy** |  |  |  |  |  |  |
| **t** | 14.11 | 15.02 | 32.55 | 18.92 | 13.59 | 16.78 |
| **p** | 0.94 | 0.92 | 0.11 | 0.6295 | 0.9708 | 0.6857 |
| **Reaction times** |  |  |  |  |  |  |
| **t** | 15.31 | 17.73 | 129.77 | 133.73 | 183.02 | 45.20 |
| **p** | 0.91 | 0.81 | $p < 10^{-5}$ | $p < 10^{-5}$ | $p < 10^{-5}$ | $p = 0.005$ |

Using similar model comparison, we also tested for an effect of the stimulus level or condition. This resulted in an AIC weight corresponding to an AIC weight of above 99% for 10 (of 12) datasets for accuracy and 10 (of 12) datasets for reaction times, showing that the stimulus manipulations were also predictive of behaviour.

For these analyses a number of trials had to be excluded for the following reasons: either the phase of respiration was not defined at the time point of interest, or the reaction time was deemed an outlier. The latter were defined as overly short (<200ms) or excessively long reaction times (defined by the median absolute deviation). Finally, few trials had to be removed as participants pressed a wrong button. Collectively this led to the exclusion of 8.6±3.4% (mean±SD) of the available trials. Overall, we included 107’432 trials from 277 participants in the analysis.

To visualize the behavioural data against the trial-wise respiratory phase (Fig. 4) we partitioned the phase angles into 6 equally-spaced bins and computed for each participant the fraction of correct responses and the median reaction time (based on the square-root transformed data, as used for linear models) for each bin. To emphasize variability over phase bins rather than participants we subtracted the bin-averaged performance (reaction time) for each participant.

### Analysis split by respiration rate

In a separate analysis we split participants into those breathing generally faster and those breathing slower. This was implemented using a median split on the average duration of the respiratory cycle. Because the number of fast and slow breathers differed between datasets, we probed the predictive power of respiration on behaviour for each group as follows: we derived the group-level AIC values by fitting linear models to the individual participant data (similar to eq. 1, but without the participant id as random effect). The individual AIC values were then summed across participants per group (Fig. 5B).

To visualize the respiratory time course for fast and slow breathers separately (Fig. 5A), we first determined for each group a prototypical respiration curve. This was obtained by finding that participant who’s inspiration and expiration durations were closest to the average of the respective groups and computing the trial-averaged shape of one respiration cycle of this participant. This cycle was then appended a few times to cover the time window to be displayed (note that this implicitly assumes that subsequent cycles are identical). This prototypical curve was then shifted so that the phase at the time of response reflects the group-average phase at this time.

To relate the trial-average respiratory phase to behaviour (Fig. 6), we implemented the following in-between participant analysis. We derived for each participant the trial-averaged phase at -2.1s prior to the response, and divided these phase values into the six equally-spaced phase bins. We then visualized the participant-wise fraction of correct responses (across all trials) and the median reaction time (across all trials) against these bins. To avoid outliers, we only included participants whose median reaction time was below 2.5s. Because many participants share a similar average phase, the effective sample size per phase bin differs. For a statistical comparison we hence contrasted the data in the two non-optimal bins (red/yellow) against the two optimal bins (blue/magenta) combined. The effect sample sizes in these bins were as follows: n=[55,69] for fast breathers and n=[28,69] participants for slow breathers.

## Results

The average duration of respiratory cycles was 3.6±0.7 s (mean±SD). As typical for spontaneous respiration, exhalation periods were longer (1.9±0.4 s) than inhalation periods (1.6±0.4 s).

### Respiration is aligned to the experimental trials

To confirm that participant’s respiration was systematically structured relative to the experimental trials, we measured the between-trial consistency of the respiratory phase using a phase-locking index. Figure 1 shows the group-level results for all trials (panel A), trials split by reaction time (B), response accuracy (C) and by time-on-the-experiment (D); the data are also shown separately for the phase of respiration relative to stimulus onset (upper panels) and the times when participants responded (lower panels). Phase locking was stronger than expected by chance (randomization tests, p<0.001; see grey line in Figure 1A) and was systematically higher around the stimulus/response period (near time 0s) compared to a few seconds earlier. We did not observe significant differences in phase locking when splitting trials by accuracy, reaction times or time-on-the-experiment (Wilcoxon sign-rank tests, at time points t=0s, all p>0.54, Z<0.61).

**Figure 1.**
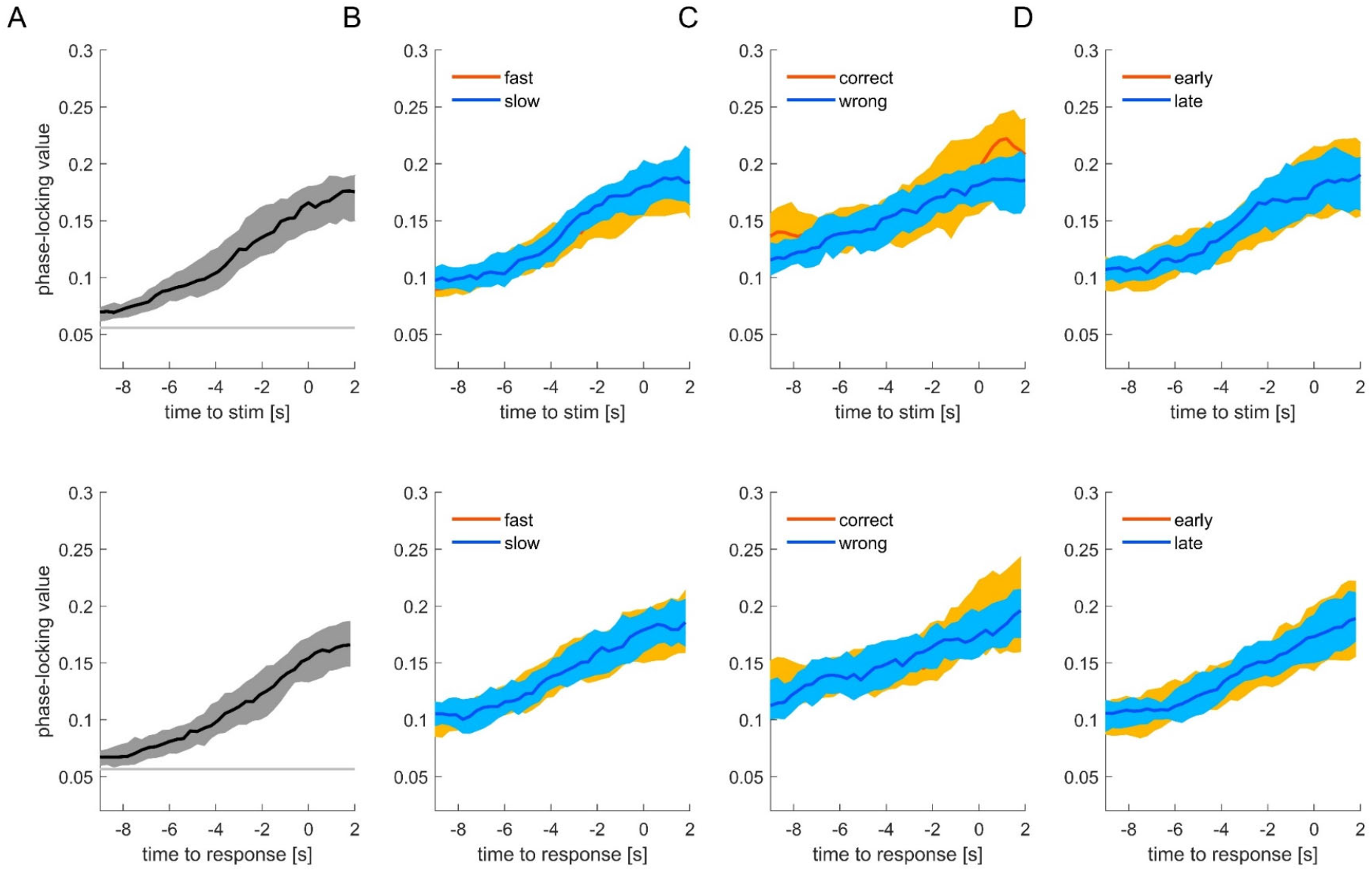
Consistency of respiratory phase across trials within participants. The figure shows the group-level phase locking values for the data aligned to stimulus onset (upper panels) or participants’ responses (lower panels). The four panels show the phase locking computed across all trials (**A**), trials split by the participant-wise median reaction times (**B**), response accuracy (**C**) and the time-on-experiment (**D**). Lines indicate the group median, shaded areas the 95% bootstrap confidence interval. The grey horizontal line in panel A shows the value corresponding to a significance of p<0.001 (randomization test; corrected for multiple tests across time points). The sample size is 277 for all panels except C, where some participants had to be excluded as they featured too few trials to estimate phase-locking separately for correct and wrong trials (n=208).

Figure 2 shows the trial-averaged respiratory phase for each participant (panel A). While Figure 1 shows that the respiratory phase is consistent across trials within each participant, the data in Figure 2 show that the trial-averaged average phase is also similar across participants. Panel B illustrates the average phase together with the phase locking strength for each participant. Phase locking is significantly stronger for respiration aligned to stimulus onset compared to response times, though the numeric differences are small (median values 0.191 vs. 0.184; Wilcoxon signed rank test, p<0.0001, Z=3.9). The between-participant consistency of the participant-wise phase angles (again measured using a phase-locking index) did not differ significantly between stimulus and response-aligned data (0.142 vs 0.108; randomization test, p=0.14), suggesting that participants respiratory phase is systematically organized around the individual trials both within and between participants.

**Figure 2.**
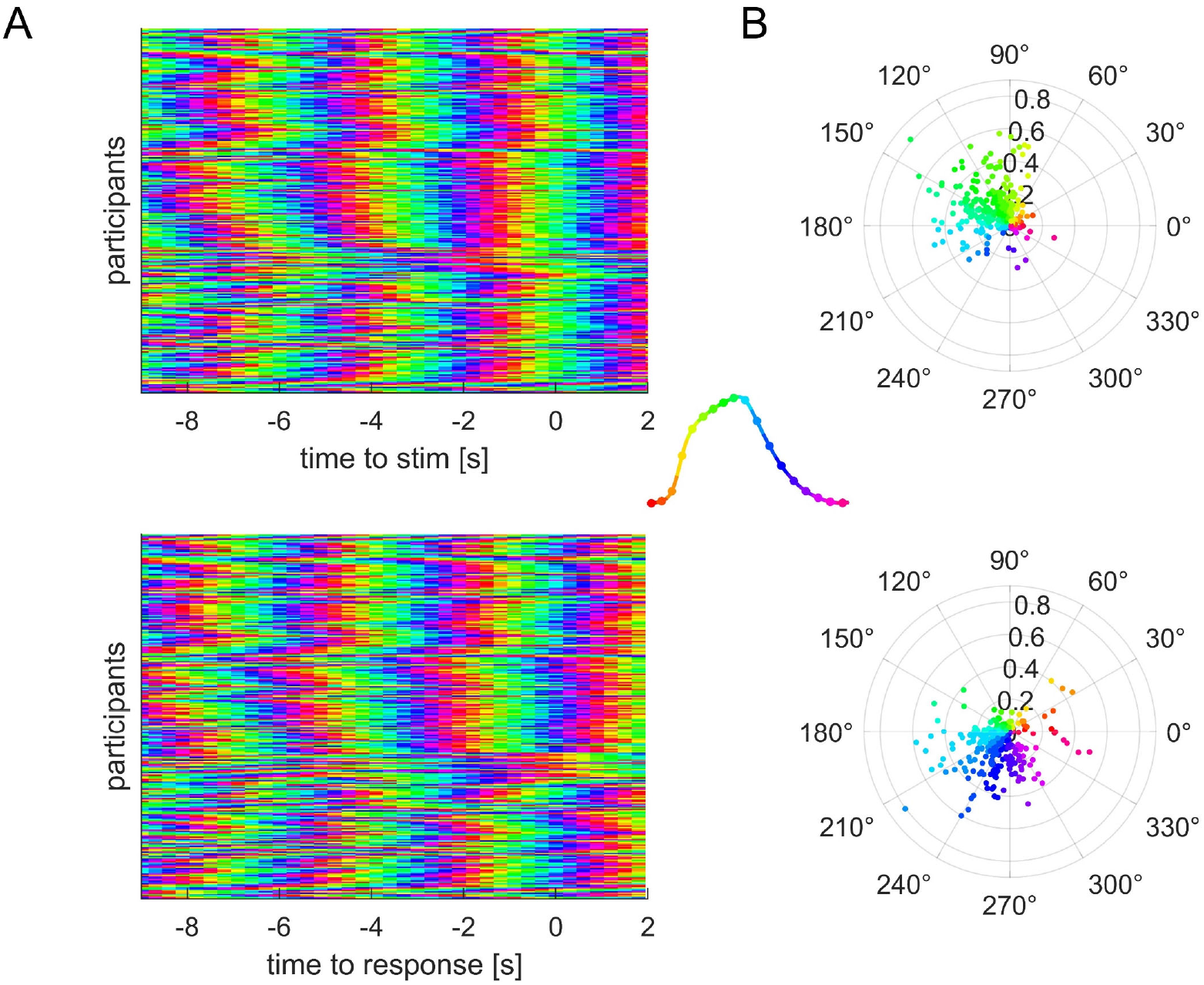
Trial-averaged respiratory phase for each participant. (**A**) Average phase for the respiratory phase derived relative to stimulus onset (top) or response times (bottom). Note how the phase becomes more similar across participants around each trial (near time zero). (**B**) Polar plot of the participant-wise average phase together with the participant-wise phase-locking value (coded as distance from the centre). Note how the population shifts from a greenish phase to blueish phase from stimulus onset to the response. The inset in the middle shows the color-coding of phase along a prototypical respiratory cycle (inhalation upwards).

### Behavioural performance is modulated along the respiratory cycle

We tested whether the respiratory phase has predictive power on the trial-wise response accuracy or reaction times. For this we fitted linear mixed models to the data from each experiment and relied on model comparison to test for an effect of respiration. Figure 3 shows the AIC differences for an effect of respiratory phase tested at different time points prior to the stimulus onset or the response (in 300ms steps). The strongest evidence for a co-modulation of behaviour with respiration was observed for the response-aligned data. Here, the evidence for a co-modulation of reaction times with respiration was very strong and the AIC weight (probability of the model including respiration to fit the data better) was above 99.999%. For 10 (of 12) datasets did we obtain an AIC weight corresponding to at least 99% in favour of an effect of respiration. For response accuracy, in contrast, only 4 datasets featured an AIC weight corresponding to a probability of above 95%. At the group level, the AIC difference was in favour of no effect of respiration. Importantly, by considering the respiratory state at various time points prior to the trial, the present data show that the peak relation between respiration and behaviour emerges around 2 seconds prior to participant’s responses. To directly confirm this, we compared the predictive power of a model relying on the phase of respiration defined at 2.1s prior to participants response (the peak time) with one relying on the phase directly at the response time: this provided clear evidence in favour of respiration defined around 2.1s prior to the response (delta AIC = 129, AIC weight of above 99.999%).

**Figure 3.**
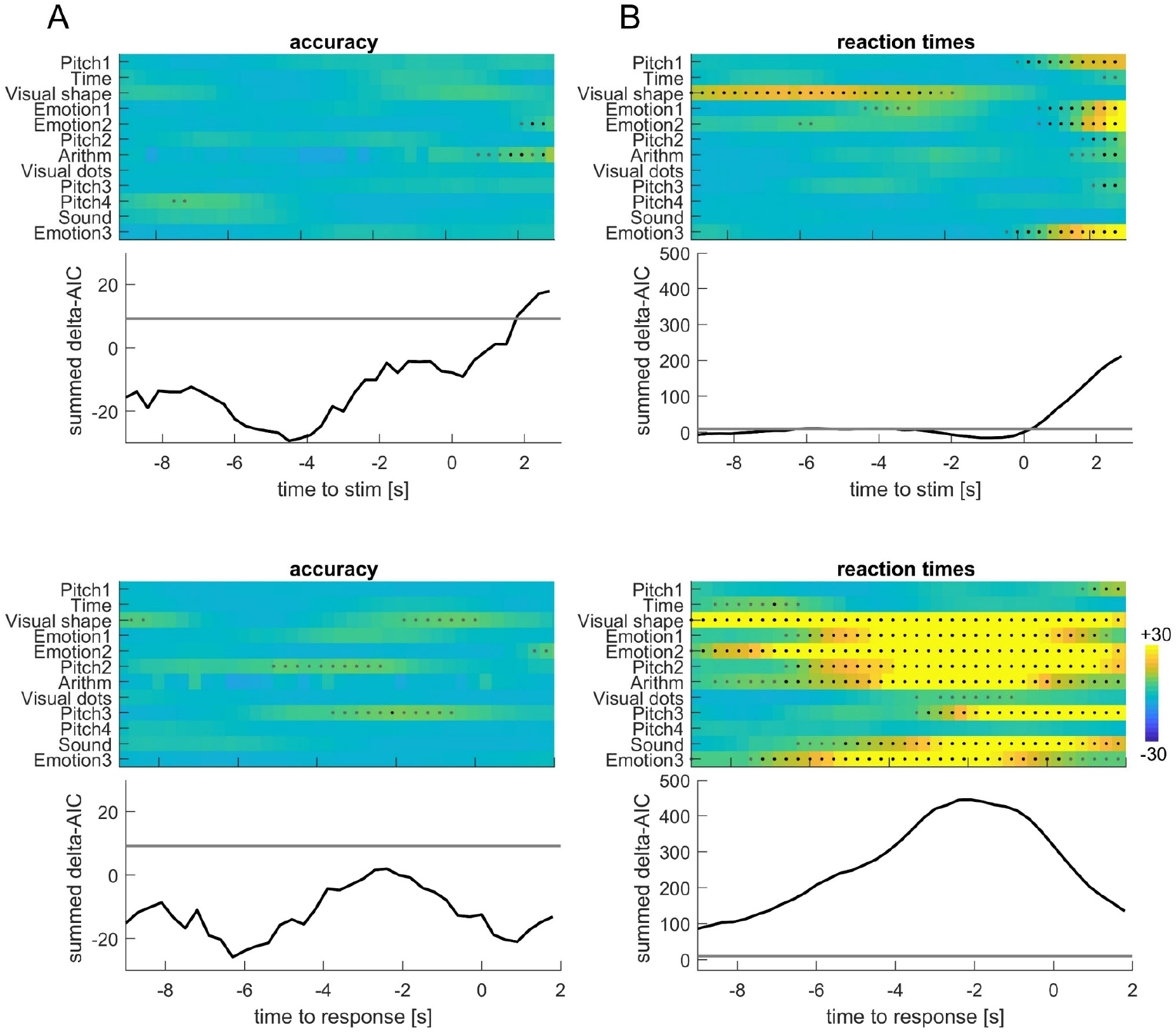
Co-modulation of behaviour and respiration. The statistical evidence in favour of a relation between respiration and trial-wise response accuracy or reaction times is shown as the AIC difference between a model including respiration and a model not-including this as factor. Positive numbers indicate evidence in favour of an effect. The upper panels show the results for respiration aligned to stimulus onset, the lower aligned to response times. The color-coded data show the outcome for each of the datasets, the graphs the summed AIC difference across all datasets. The grey (black) dots indicate AIC differences corresponding to an AIC weight of 95% (99%) in favour of an effect of respiration.

Figure 4 illustrates this co-modulation by binning the trial-wise respiratory phase into 6 bins (defined 2.1 s prior to the response); for each bin we calculated the average reaction time and the fraction of correct responses per participant. The fraction of correct responses is highest around the blue phase, where reaction times are shortest, while the opposite is observed near the yellow/red phase. This combined co-modulation of accuracy and reaction times with respiration is shown in panel C.

**Figure 4.**
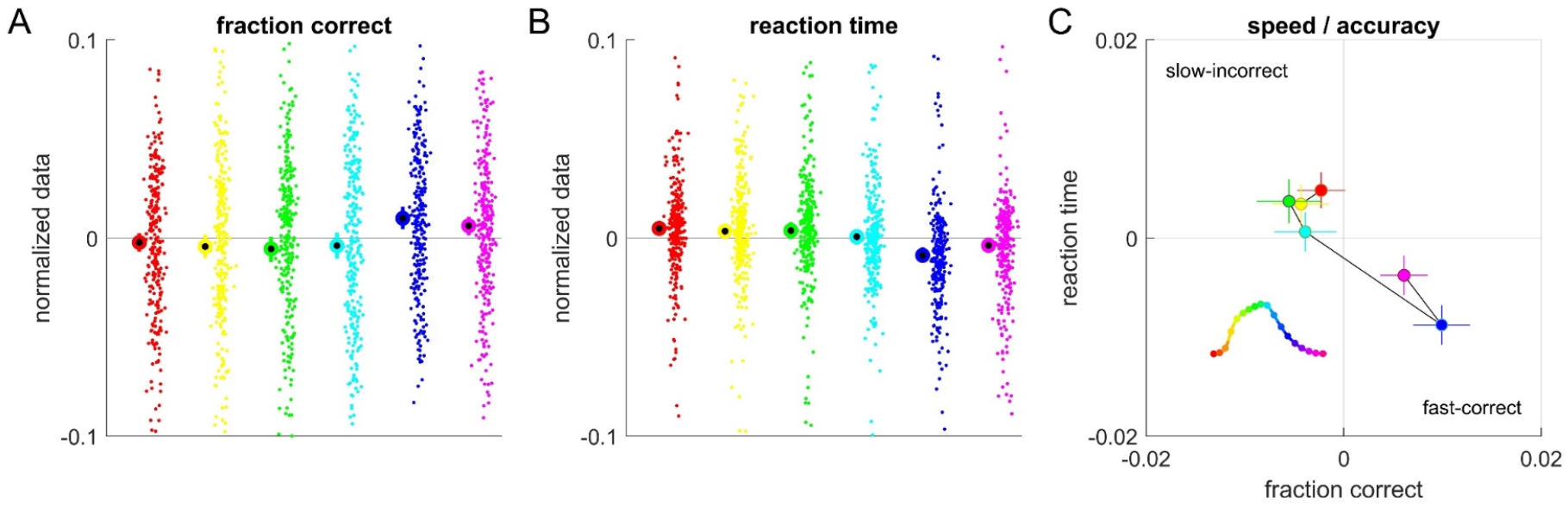
Behavioural data vs. respiratory phase. (**A**) Fraction of correct responses for trials featuring either of six (binned) respiratory phase angles. Dots indicate individual participants, thick circles the bin-average. The data were mean-normalized for each participant for better visualization. **(B)** Reaction times for each respiration phase bin. The single trial reaction times were square-root transformed and mean-normalized for each participant for visualization. (**C**) Group-level mean and s.e.m. of fraction of correct responses and reaction times for each phase bin. The respiratory phase was derived 2.1s prior to the response (corresponding to the peak in Fig. 3B).

As a further statistical test of the predictive power of respiration on behaviour, we probed whether the strength (vector-length) of the respiratory predictors for the actual data was stronger compared to surrogate data. The results (Table 1) corroborate that the respiratory predictors are highly significant, in particular for reaction times and the response-aligned respiratory phase.

### No influence of overall respiration rate

Given the inter-individual differences in average respiration rate (cycle durations: median 3.5s, range 2.1s and 6.3s), it is possible that the statistical co-modulation of behaviour and respiration is expressed differently for participants breathing generally faster and slower. For example, the duration of the individual respiratory cycle constrains how strongly respiration can be aligned to the sequence of individual trials, as longer respiratory cycles (e.g. around 6s) are more difficult to align to individual trials on the time scale of 2-3 seconds. Indeed, the average respiratory rate was significantly anticorrelated with the phase locking value (Spearman rank correlation, r=-0.57, p<10^−10^, 99% confidence interval [-0.69 -0.43]). As a result, the individual respiratory rate may affect how well between-trial variations in respiratory phase can explain between-trial variations in behaviour. To investigate this, we split the participants into those generally breathing fast or lower (median cycle durations 3.1 and 3.9s respectively).

To visualize the respiratory traces for each group we first derived the prototypical respiratory trace (trial-and participant-averaged) for each group and aligned this to the group-averaged phase at the time of response (Fig. 5; top). This reveals that both groups exhibit a similar average respiratory phase in the period prior to the response. We confirmed that for both groups there is indeed a significant co-modulation of the trial-to-trial fluctuation in reaction times and respiratory phase (Fig. 5B; peak delta-AIC values: 229 and 898; corresponding to a model probability of above 99.999%). Figure 5C shows the behavioural data for individual respiratory phase bins, illustrating that for both groups reaction times are fastest around the same phase.

**Figure 5.**
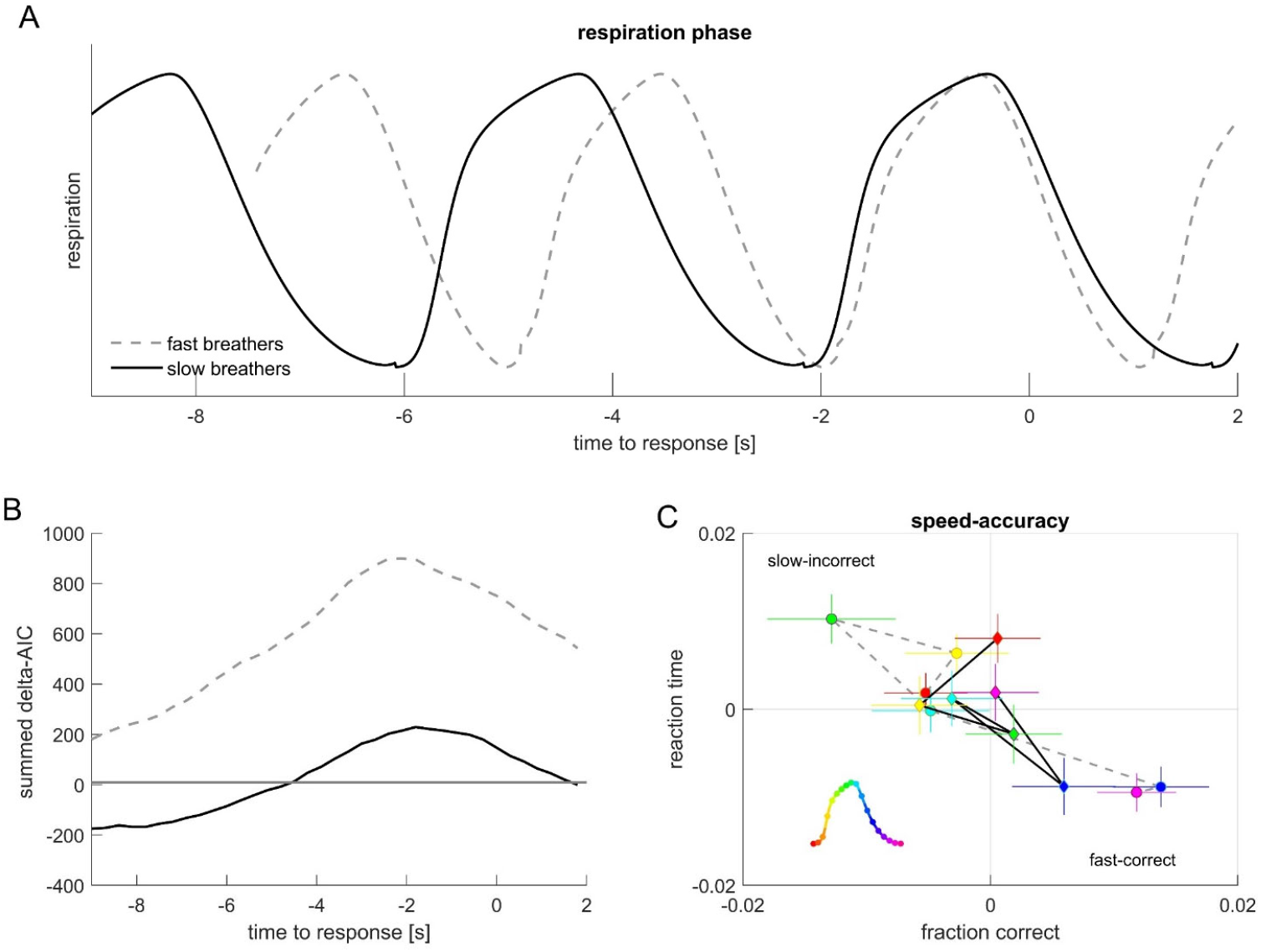
Results split by respiratory rate. For this split the sample into fast and slow breathers (median split). **(A)** Prototypical respiratory phase of each group, aligned to exhibit the response group-average phase at the time of response. Despite the different cycle durations both groups exhibit a similar transition from exhalation to inhalation around 2 s prior to the response. (**B**) Statistical evidence for an effect of respiration on reaction times, shown as group-averaged AIC values. Positive numbers indicate evidence in favour of an effect of respiration. (**C**) Group-level mean and s.e.m. of fraction of correct responses and reaction times for each respiratory phase bin (with phase derived 2.1s prior to the response time).

### Average respiratory phase is also predictive of behaviour

The above shows that participants tend to align their average respiratory phase to the experimental trials, but also that the trial-wise respiratory phase is predictive of behaviour. This suggests that participants who tend to align their respiration with the ‘optimal’ phase to the trials may on average perform better or respond faster than participants whose average phase is non-optimal. To test whether this is indeed the case, Figure 6 shows the trial-averaged behavioural data for the participant sample split by the trial-average respiratory phase. While this shows great heterogeneity in the behavioural performance across participants, the figure also suggests a difference in reaction times between those individuals aligning with the optimal phase (blue/magenta) and to those aligning with a non-optimal phase (red/yellow). Because the number of participants per phase bin differs, we restricted the statistical analysis to the comparison of reaction times between optimal and non-optimal aligners: This revealed a significant difference in the median reaction times regardless of respiration rate (separate Spearman rank-sum tests; fast breathers: p<10^−3^, Z=3.52; slow breathers: p=0.0015, Z=3.17).

**Figure 6.**
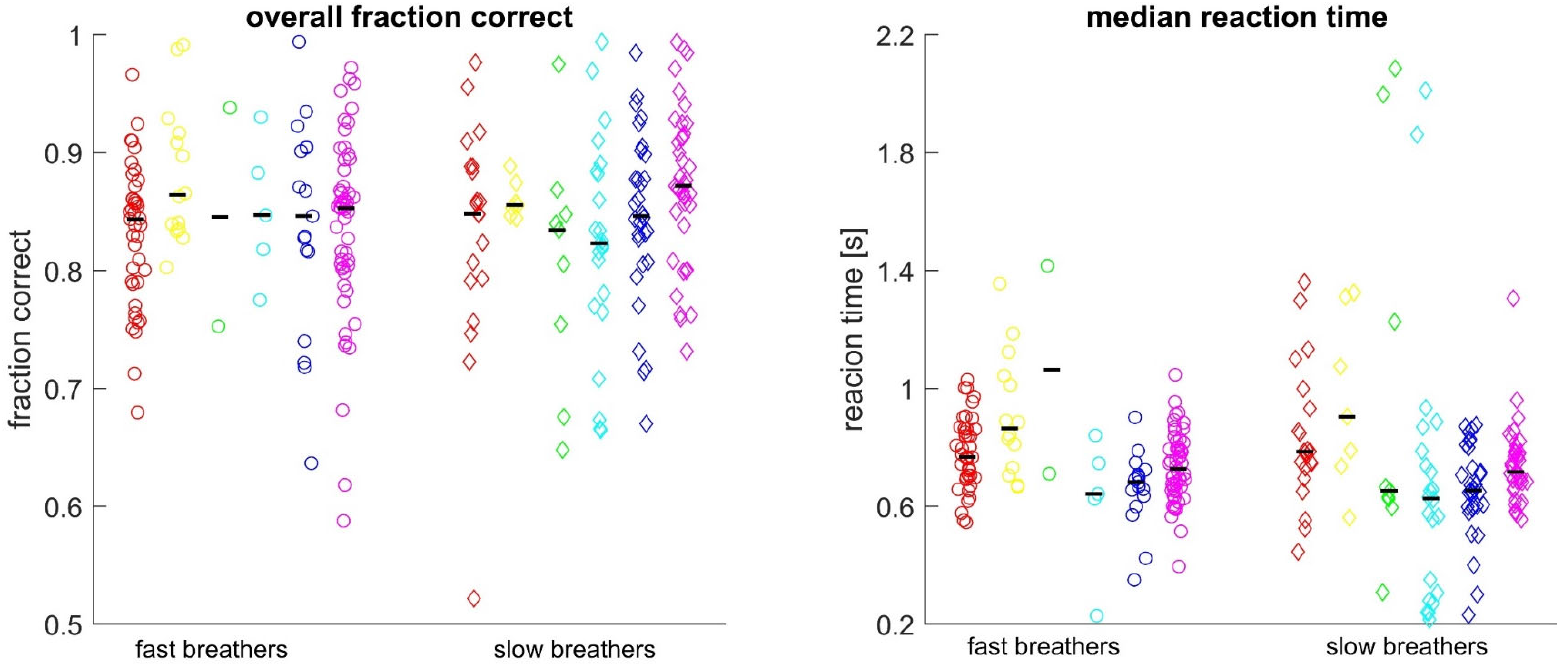
Trial-average respiratory phase is predictive of overall behaviour across participants. **(A)** Fraction of correct responses across all trials, **(B)** median reaction time (in seconds, not normalized) for each participant against their average respiratory phase (derived at -2.1 s prior to the response). Note that the number of data points per bin differs. Symbols indicate individual participants, black lines the median within each bin. For statistical analysis we compared the reaction time in the optimal bins (blue/magenta) against those in the non-optimal bins (red/yellow) in a between participant analysis.

## Discussion

Our results show that participants tend to align their respiration to the expected timing of a sequence of experimental trials. At the same time, the respiratory phase holds significant predictive power on behavioural performance both across trials for an individual and across the sample of participants. This relation of respiration and behaviour emerges regardless of whether an individual generally breathes faster or slower and is strongest when considering the respiratory phase around two seconds prior to the individual responses. Our results demonstrate these results across a large sample of data from different typical sensory-cognitive tasks, confirming the robustness and ubiquity of this relation between respiration and behaviour.

### Respiratory phase displays consistency and variability relative to experiment trials

It is known that humans tend to breath in a structured manner around expected events, such as during sports, during conversation and also in a laboratory task [25, 27, 28, 39, 40].This alignment of respiration to such events manifests is a consistency of the respiratory phase across trials. In the present data this phase-locking of respiration was comparable when computed for the stimulus- and the response-aligned data. This is not surprising given that for most paradigms studied here the duration of stimulus presentation was short and reaction times between 0.51 and 1.36 s (the time between stimulus onset and response; 10^th^ and 90^th^ percentile across all trials). This makes a differential alignment of respiration to stimulus onset and response times practically difficult, given that the typical respiratory cycle duration was around 3.4s (overall median). In a previous study we concluded that respiration was significantly stronger phase locked to the response [27]; however, also there were the numerical differences small. Given the slow duration of individual respiratory cycles compared to the faster sequence of experiment trials, it is inherently difficult to determine statistically whether respiration was aligned actively and selectively to either an expected stimulus or to the planned motor actions. Consistently across studies, however, we see the average tendency of the respiratory phase is to inhale around stimulus presentation and to exhale around responses.

Besides this alignment, the individual respiratory phase is predictive of behaviour. This was visible for individual participants in the trial-to-trial data and was stronger for the respiratory phase measured relative to the response time rather than to the stimulus onset. This implies that the trial-to-trial variability of the difference in respiratory phase between stimulus onset and response time is sufficient to account for their difference in predictive power against behaviour. Indeed, the overall median reaction time (0.73s) corresponds to half the median duration of inhalation period (1.48s), which is sufficient to shift the respiratory phase from a phase angle related to faster response to one related to slower responses (c.f. Fig. 4). At the same time, we found that also the trial-averaged respiratory phase is predictive of the between-participant variations in reaction times: participants who align ‘optimally’ to the paradigm tended to respond faster than those how aligned ‘non-optimally’. Hence, respiratory phase has a profound and systematic relation to the speed of responses.

### Time-lagged relation of respiration to behaviour

A central result is that the predictive power of respiration on behaviour peaks about two seconds prior to the response, independently of how fast an individual breathes. This result departs from previous studies, which mostly investigated the state of respiration at a single time point and did not probe for a systematic time lag between respiration and behaviour.

This time lag provides clues about the putative mechanisms underlying the co-modulation of respiration and behaviour. A short-latency co-modulation may be well explained by direct neural feedback about respiratory muscle movements or the resulting airflow [5-8]. In contrast, a time-lagged effect also leaves the possibility that respiration-related changes in heart rate or blood oxygenation contribute. While it takes several seconds for oxygen to reach the brain from the lungs, the time scale of 2 seconds is also sufficient to for example detect changes in the instantaneous heart rate. The heart rate and respiratory phase are directly and indirectly coupled by multiple mechanisms and common inputs from the autonomic nervous system [41]. Information about the timing of the heart beat is transported by the vagus nerve and is available to interoceptive brain regions [42-45]. Similar as for respiration, studies show that the relative timing of stimuli to the heartbeat shapes how they are perceived or memorized [46-49]. Furthermore, respiration is also known to modulate cortical signatures of neural excitability and arousal [13, 24], possibly by modulating activity of the locus coeruleus [5, 50]. To better understand the pathways by which a time-lagged influence of respiration on sensory-cognitive processes and motor actions emerges, future work needs to dissociate the multiple bodily and neuro-physiological signals and their relation to behaviour [4].

### Does respiration relate to both reaction times and response accuracy?

The literature on a co-modulation of respiration and performance in neuro-cognitive tasks features both effects for reaction times and response accuracy. As we recently showed [27], changes in response accuracy along the respiratory cycle are typically between 1% to 5%, and many studies failed to find significant changes in accuracy. In contrast, changes in reaction times are reported more frequently and amount to effect sizes comparable to changes in induced by brain stimulation or classical cognitive interference effects [27]. In the present data the evidence for a relation between respiration and behaviour was generally stronger for reaction times compared to accuracy. Only for 4 out of 12 datasets did we find a significant predictive power of respiration on response correctness.

While this may suggest that response speed is generally more influenced by respiration than the accuracy of individual responses, one cannot rule out that differences in the statistical sensitivity of the respective linear models contribute to this result. Given the continuous and discrete nature of the trial-wise reaction times and accuracy values, we relied on different statistical models for these data; this may at least theoretically result in a differential sensitivity. Indeed, the group-level averages Fig. 4) reveal a visible modulation in both aspects of behaviour. However, in a previous study we relied on a different statistical approach (see also below), which did not require different models for reaction times and accuracy [27]. Still, we found more prominent effects for reaction times. All in all, this calls for future studies to more finely dissociate effects of respiration on the quality of sensory representations that may shape response accuracy and more peripheral effects such as muscle tone or brain-muscle communication [13, 23].

### Comparison to previous work

In a previous study we had analysed some of the present data using slightly different procedures [27]. While the conclusions drawn previously are supported by the present report, there are a number of technical details that differ between reports that are worthwhile mentioning. First, compared to previous work, we used a revised pre-processing pipeline which assigned a defined respiratory state to a larger fraction of trials, hence losing less of the data. Second, the previous statistical analysis relied on phase-binned data, while in the present study we model the single-trial data against respiration. The latter is particularly relevant when also including additional variables, such as brain activity [1], but also helps avoiding potential biases induced by data binning in general [51, 52]. And third, in the previous study we used both the overall state of respiration (inspiration /expiration) and the phase within each of these states as separate predictors. While splitting state and phase seems conceptually appropriate given that inspiration and expiration are physiologically and mechanistically distinct, any downstream influence of these can be time-shifted relative to these by an arbitrary amount and hence this dichotomous interpretation may not be relevant for the interpretation of such effects.

## Conclusion

Making repeated judgements or motor actions comes with trial-to-trial variability in the accuracy and speed of these. Trial to trial variations in respiratory phase do explain some of this variability and our results show that this co-modulation follows a time lag. At the same time respiration is also systematically aligned to expected events, and the phase of alignment is predictive of behaviour across participants. This suggests a profound co-modulation of respiration and perceptual and cognitive performance, which may form the basis for better understanding how conscious respiration and breath-practices can improve brain function or mood, in sports, therapeutic applications or every-day scenarios [53, 54].

## Acknowledgements

We thank Alex Wecker for developing the respiration sensors and Michelle Johannknecht for data collection and intellectual contributions to this line of work.

## Notes

### Competing Interest Statement

The authors have declared no competing interest.

